# Blindness reveals that Heschl’s gyrus folding is not altered by auditory experience

**DOI:** 10.64898/2026.04.30.721986

**Authors:** Amy Poole, Kelly Chang, Feiyi Wang, Ione Fine, Woon Ju Park

**Affiliations:** Department of Radiology, University of Minnesota, 55455; Department of Psychology, University of Washington, 98195; Department of Psychological & Brain Sciences, Boston University, 02215; School of Psychology, Georgia Institute of Technology, 30332

## Abstract

Heschl’s gyrus (HG), which contains the primary auditory cortex, shows marked individual variability in its folding pattern, ranging from a single gyrus to partial or complete duplication. Greater HG duplication has been reported in expert musicians, often interpreted as evidence that auditory experience can shape cortical morphology. However, these structural differences might alternatively indicate a bias for musical careers in individuals whose anatomical predispositions facilitate expertise. Here, we examined HG morphology in blind individuals—a population with extensive auditory experience but without selection based on auditory ability. T1-weighted MRI data from 100 human participants (48 females, 42 males, 10 unknown) across blind and sighted groups were analyzed. HG was manually defined in each hemisphere, and folding was measured using both categorical morphology classification and continuous surface-based metrics. Across all analyses, blindness did not increase HG folding. These results suggest that the morphology of HG is largely predetermined.

**Significance statement:** Increased anatomical folding in the auditory cortex has been reported in professional musicians. Is this structural variability due to experience-dependent plasticity, or is it that individuals with increased anatomical folding are more likely to become musicians? We examined Heschl’s gyrus (HG), which contains the primary auditory cortex, in blind individuals who rely heavily on auditory input. Despite extensive auditory experience, blindness did not alter HG folding. This finding suggests that the morphology of HG is not strongly influenced by auditory experience.

## Introduction

Heschl’s Gyrus (HG), a region on the superior surface of the temporal lobe containing the primary auditory cortex (Abdul-Kareem & Sluming, 2008; Da Costa et al., 2011; Henderson et al., 2023a; Mangold & Das, 2024; Penhune et al., 1996; Rademacher et al., 1993), is known for its high morphological variability among individuals and hemispheres. HG’s morphology is typically classified into three types: single gyrus, partial duplication (also known as a common stem duplication), or complete duplication (also known as a complete posterior duplication; **Figure 1**). Traditionally, these categories have been defined through manual labeling, which remains a gold standard for identifying and localizing the region (Abdul-Kareem & Sluming, 2008; Henderson et al., 2023b; Kasai et al., 2003; Leonard et al., 1993; Marie et al., 2015; Schneider et al., 2009), despite recent efforts to automate the process (Dalboni da Rocha et al., 2023).

**Figure 1.**
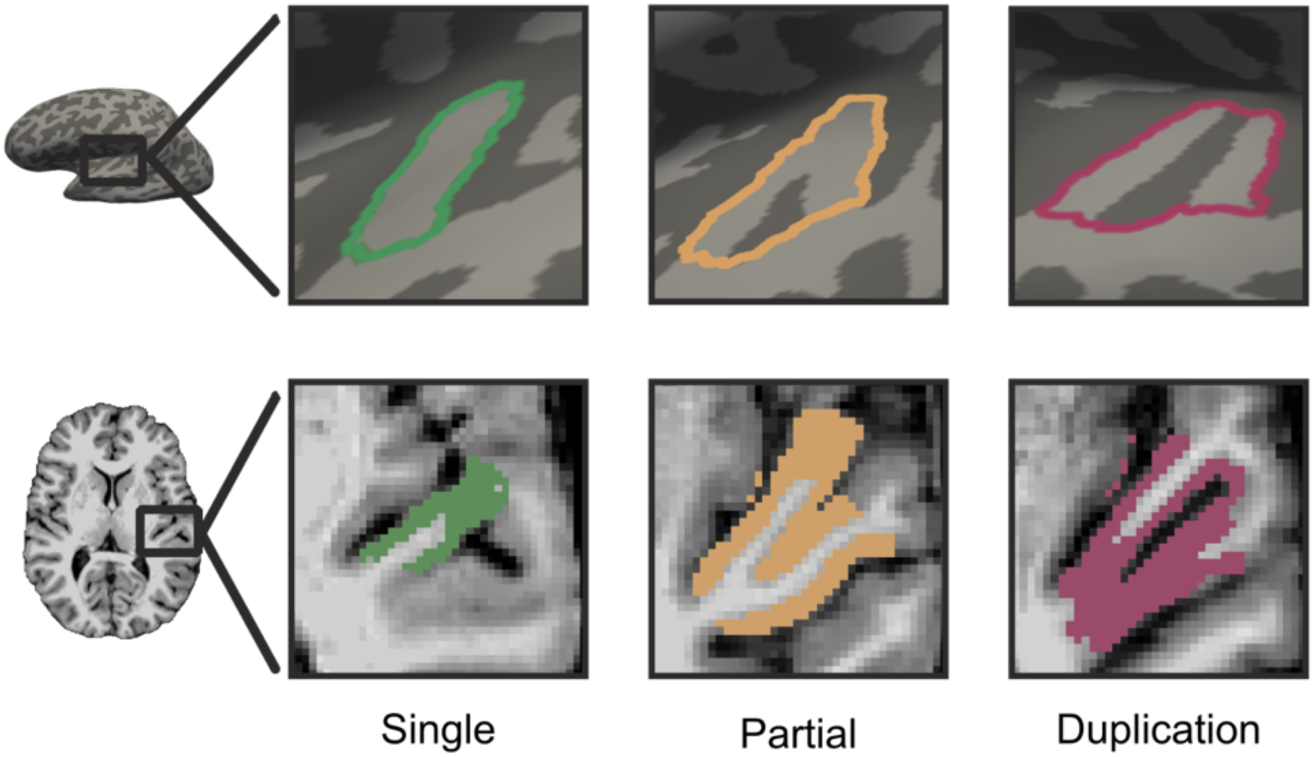
Heschl’s gyrus (HG) folding categories in the left hemisphere in both the inflated cortical surface (top) and the volume (bottom), zoomed in on HG to show the differences between each category. Single (outlined in green) on the left is one gyrus. Partial (outlined in tan) has a sulcus splitting part of the gyrus. Duplication (outlined in magenta) on the right has two separate gyri.

The structure of HG has been linked to individual differences in acoustic performance across both music and speech (Herholz & Zatorre, 2012; Zatorre, 2013). Up to 90% of musicians who practice music approximately 17 hours per week show HG duplications in one or both hemispheres, a prevalence significantly higher than in the general population (Benner et al., 2017). Relatedly, expert phoneticians are more likely than controls to exhibit left-hemisphere duplications (Golestani et al., 2011). These findings suggest that increased HG folding is associated with auditory expertise. However, an important question remains: does heightened auditory demand drive changes in HG folding, or do individuals with greater folding from birth have a predisposition toward careers that require auditory expertise?

Cortical folding is generally thought to develop prenatally; patterns of cortical folding are highly heritable (Schmitt et al., 2021) and predictable within species (Van Essen et al., 2019). Maturation across the lifespan follows a stereotypical pattern in which gyrification peaks during childhood (Li et al., 2014; Raznahan et al., 2011) and linearly decreases over time as one ages (Hogstrom et al., 2013; Madan, 2021). Unlike other measures of cortical structure shown to be shaped by experience, such as the gray matter volume or cortical thickness in taxi drivers (Lövdén et al., 2013; Maguire et al., 2000; Thomas & Baker, 2013), the morphology of folding within a cortical area has generally been considered relatively resistant to experience-dependent plasticity. However, it remains possible that folding in areas like HG, which show large amounts of variability across individuals, might depend on experience.

In this study, we examine this question by comparing HG folding across sighted and blind individuals. Blind individuals rely heavily on sound to understand and navigate the world; like musicians or phoneticians, they show pronounced expertise in relevant auditory tasks (Gougoux et al., 2004; F. Jiang et al., 2014; Röder et al., 2001; Stevens & Weaver, 2005). Unlike musicians and phoneticians, being blind is not a “choice” that might be influenced by a biological predisposition to become an auditory expert. Any differences in HG morphology among blind individuals must be due to auditory experience.

We compared HG duplication and gyrification across four groups of participants: anophthalmic individuals whose eyes failed to develop and thus received no signals from the eye to the brain during development in utero; early blind individuals who lost their vision at birth or very soon afterward; late blind individuals who had normal vision until adulthood; and sighted controls who have normal vision. HG folding was measured in each hemisphere from T1-weighted MRI images, first by manually categorizing the morphology type, and then by mathematically characterizing the Gyrification and Curvedness Indices. Across all measures, we find no evidence that blindness and the extensive auditory experience that accompanies it lead to increased folding within HG. Additional exploratory analyses that assessed cortical thickness, surface area, and gray matter volume also failed to find structural differences between blind and sighted groups.

## Materials and Methods

### Participants

One hundred participants’ structural T1-weighted MRI images were drawn from five studies across three sites (University of Washington, University of Pennsylvania, Oxford University; Aguirre et al., 2016; Bridge et al., 2009; Jiang et al., 2016). Six participants had anophthalmia (ANO), meaning that they were born without eyes and have been blind since birth. Forty-eight acquired blindness before the age of 4 and were labeled as early blind (EB). Eighteen individuals became blind after the age of 4 and were labeled as late blind (LB). The remaining 28 participants had normal or corrected-to-normal vision (sighted controls; SC). See **Table 1** for demographic information.

**Table 1.**
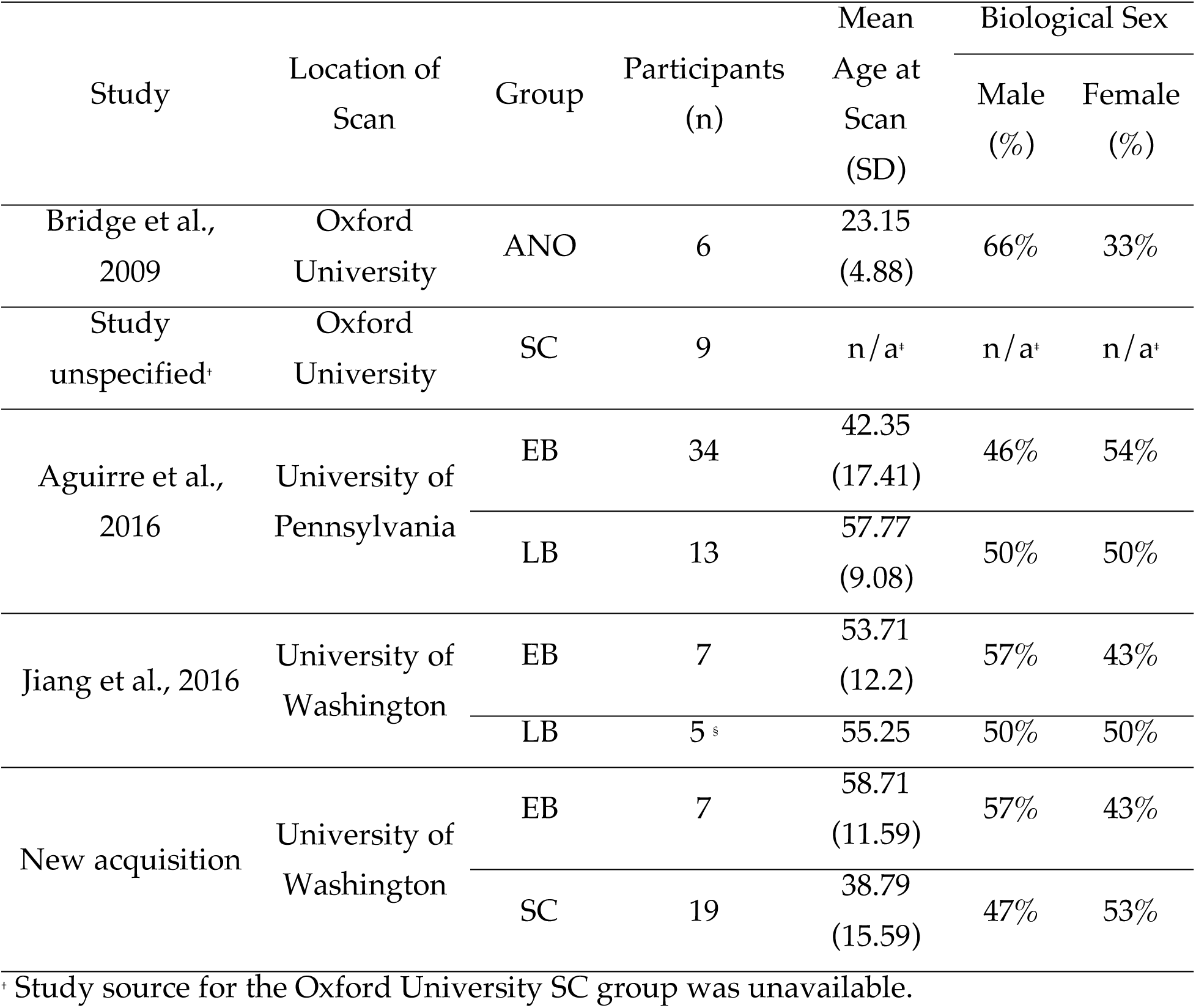
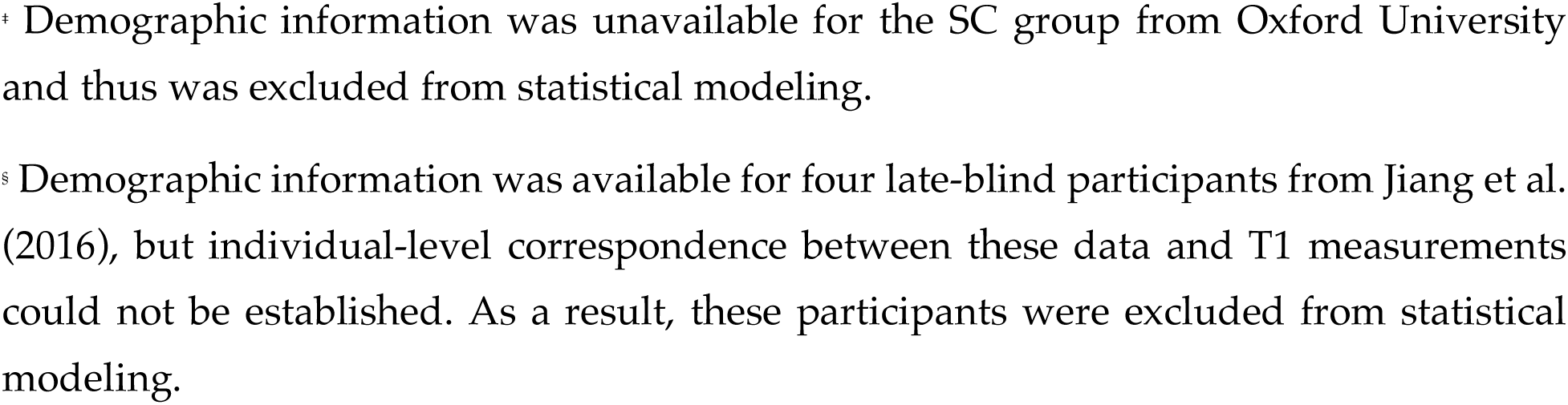
Demographic information by dataset. Details include the location where each dataset was acquired, the sample size by blindness groups, and biological sex. The mean age for each study dataset is shown with the standard deviation in parentheses. There are 14 individuals of unknown age: nine sighted controls from Oxford, and five late blind from the University of Washington (F. Jiang et al., 2016).

| Study | Location of Scan | Group | Participants (n) | Mean Age at Scan (SD) | Biological Sex |  |
| --- | --- | --- | --- | --- | --- | --- |
|  |  |  |  |  | Male (%) | Female (%) |
| Bridge et al., 2009 | Oxford University | ANO | 6 | 23.15 (4.88) | 66% | 33% |
| Study unspecified<sup>†</sup> | Oxford University | SC | 9 | n/a<sup>†</sup> | n/a<sup>†</sup> | n/a<sup>†</sup> |
| Aguirre et al., 2016 | University of Pennsylvania | EB | 34 | 42.35 (17.41) | 46% | 54% |
|  |  | LB | 13 | 57.77 (9.08) | 50% | 50% |
| Jiang et al., 2016 | University of Washington | EB | 7 | 53.71 (12.2) | 57% | 43% |
|  |  | LB | 5 <sup>s</sup> | 55.25 | 50% | 50% |
| New acquisition | University of Washington | EB | 7 | 58.71 (11.59) | 57% | 43% |
|  |  | SC | 19 | 38.79 (15.59) | 47% | 53% |

<sup>†</sup> Study source for the Oxford University SC group was unavailable.
\* Demographic information was unavailable for the SC group from Oxford University and thus was excluded from statistical modeling.
§ Demographic information was available for four late-blind participants from Jiang et al. (2016), but individual-level correspondence between these data and T1 measurements could not be established. As a result, these participants were excluded from statistical modeling.

### MRI Data

At the University of Washington, earlier images from Jiang et al. (2016) were collected at the Diagnostics Imaging Sciences Center using a 3T Philips scanner. At the University of Pennsylvania, images from Aguirre et al. (2016) were acquired using a 3T Siemens Trio scanner with an 8-channel head coil. Finally, at the University of Oxford, images from Bridge et al. (2009) and additional data from an unspecified study were acquired at the Oxford Centre for Clinical Magnetic Resonance Research using a Siemens 3T Trio scanner with a 12-channel head coil. T1-weighted (T1w) MPRAGE acquisition parameters for the earlier University of Washington, University of Pennsylvania, and Oxford datasets have been previously reported (Aguirre et al., 2016; Bridge et al., 2009; F. Jiang et al., 2016), with all scans acquired at 1×1×1 mm isotropic resolution. The most recent University of Washington dataset was collected at the Center for Human Neuroscience on a Siemens 3T Prisma scanner with a 64-channel head coil using a 3D volumetric navigator-guided multiecho T1w sequence at 0.8×0.8×0.8 mm isotropic resolution (FOV = 166.4×240×256 mm; Tisdall et al., 2012).

### Preprocessing

T1w images were preprocessed with FreeSurfer-7.4.1 using the standard recon-all pipeline. This pipeline produced reconstructed 3D cortical surfaces for each hemisphere that were used to manually define HG.

### Region of Interest (ROI) Segmentation

HG ROIs were manually drawn on the cortical surface for each hemisphere of every participant. Each HG ROI was categorized as a *single*, *partial*, or *complete* duplication (see **Figure 1** for examples). When identifying and categorizing ROIs, we followed protocols from previous studies (Abdul-Kareem & Sluming, 2008; Henderson et al., 2023a). Two independent raters (A.P. and K.C.) categorized each ROI to ensure reliability. Additionally, volume slices of the ROIs were inspected to verify the accuracy of segmentation and classification. The criteria used to define each HG morphology category are described below.

### Visual Categorization of HG Folding

Manual labeling remains the gold standard method for determining HG morphology, but the exact methods and classification criteria vary widely across studies (Da Costa et al., 2011; Henderson et al., 2023b; Marie et al., 2015; Smith et al., 2011; Warrier et al., 2009; Wong et al., 2008). A key source of variability concerns how partial duplications are defined. Most consider a partial duplication to have a sulcus intermediate (SI) present of any size while still having a common stem (Abdul-Kareem & Sluming, 2008; Da Costa et al., 2011; Dalboni da Rocha et al., 2023). However, others consider HGs with a longer common stem to be a single (Benner et al., 2017; Marie et al., 2015) or those with a smaller common stem to be a complete duplication (Wong et al., 2008).

To account for these differences, we used three different approaches to characterize HG morphology as outlined below. We first adopted two established approaches (Da Costa et al., 2016; Marie et al., 2015) that varied in how they define partial duplication. We also developed a novel ranking method that explicitly accounts for the variability in the location of the gyral split along the common stem, allowing for a more graded characterization of HG morphology. These approaches were applied to evaluate HG morphology across participant groups separately by hemisphere. Lastly, the Da Costa et al. approach was used to examine the HG morphology jointly across hemispheres.

#### Da Costa et al. (2011) method

Following Da Costa et al. (2011), the HG morphology in each hemisphere was classified into one of three categories: single, partial duplication, and complete duplication (**Figure 2A**). A single HG was defined as one gyrus. A partially duplicated HG was defined as a single gyrus that separates to create an SI while maintaining a common stem. The common stem is typically medial; however, a split on both sides is often observed, giving HG the appearance of an “H” or a bowtie (Leonard et al., 1998) as compared to a typical “V” shape. A complete duplication included a second gyrus (G2) that was fully separated from the first gyrus (G1) by the SI, where neither shared a common stem.

**Figure 2.**
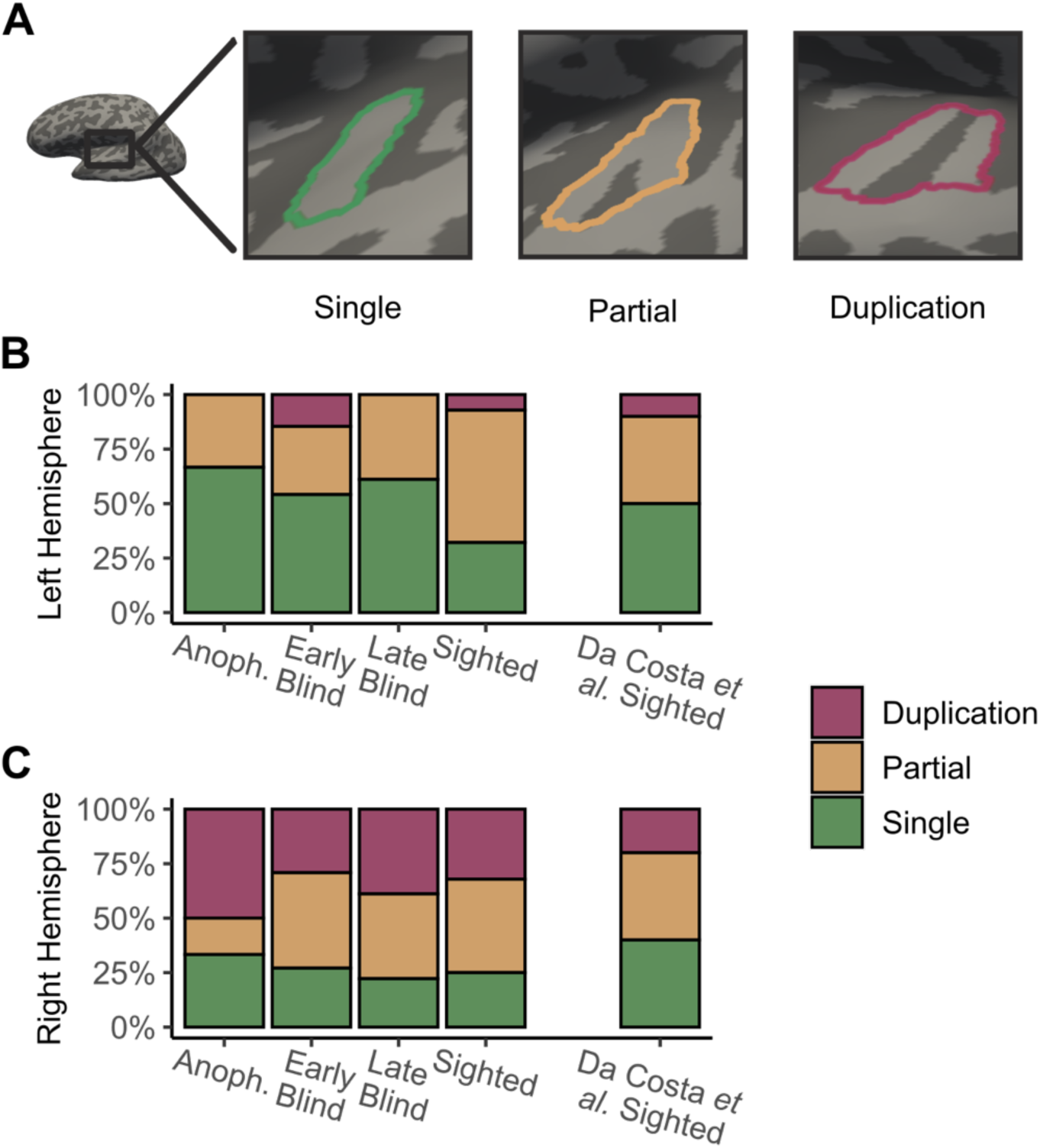
Heschl’s gyrus (HG) folding across blindness groups categorized with the Da Costa et al. method. (A) Inflated left hemisphere surface showing HG categories: single (green; one gyrus), partial (tan; sulcus partially splitting the gyrus), and duplication (magenta; two separate gyri). (B, C) Percentage of each category across groups in the left (B) and right (C) hemispheres. Groups (left to right): anophthalmia (Anoph.), early blind, late blind, sighted controls (this study), and Da Costa et al.’s sighted controls.

#### Marie et al. (2015) method

The method from Marie et al. (2015) also categorizes HG into three morphological types: single, partial duplication, and complete duplication. However, their partial duplication definition differs from that of Da Costa et al. (2011). In this approach, a duplication is considered partial only if the separation extends at least one-third the length of HG from the lateral edge. Shallower separations were categorized as single (**Figure 3A**).

**Figure 3.**
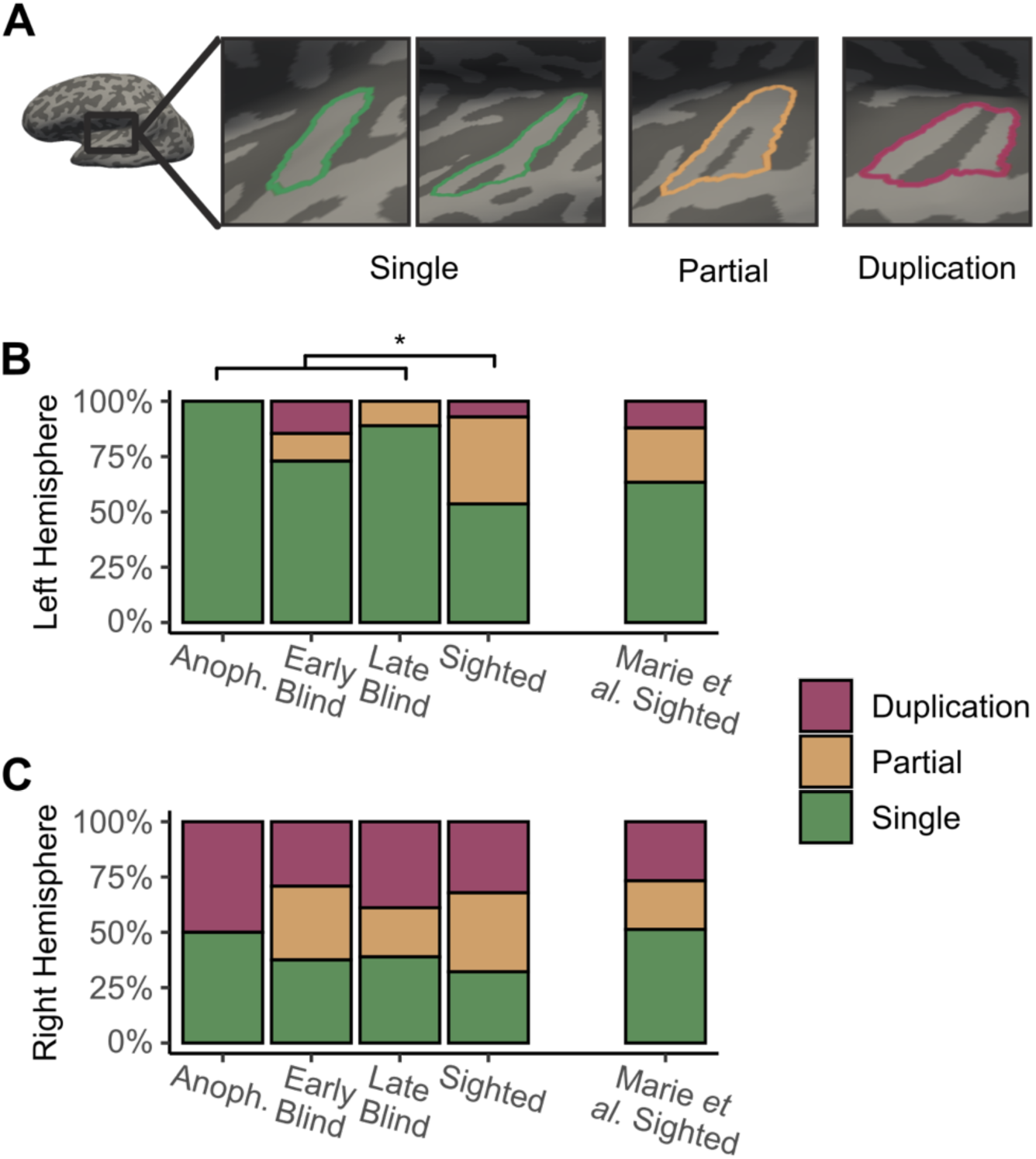
Heschl’s gyrus (HG) folding across blindness groups categorized with the Marie et al. method. (A) Inflated left hemisphere surface illustrating HG categories: single (green; includes duplications <1/3 of HG length), partial (tan; sulcus >1/3 length), and complete duplication (magenta; two separate gyri). (B, C) Percentage of each category across groups in the left (B) and right (C) hemispheres. Groups (left to right): anophthalmia (anoph.), early blind, late blind, sighted controls (this study), and Marie et al.’s sighted controls. Blind groups show a higher prevalence of singles and a lower prevalence of partial duplications in the left hemisphere compared to sighted controls (p = .01).

#### Graded duplication ranking

We developed a ranking method based on the length of the common stem (**Figure 4A**). Previous literature, particularly the two methods already mentioned, differed in how they classified partially duplicated HGs, which led to inconsistent classifications. To address this, our method extends previous work (Golestani et al., 2011) by treating the degree of duplication as a quasi-continuous variable. HGs were assigned a value from 1 to 2, where 1 represents a single gyrus and 2 represents a complete duplication. Partially duplicated HGs were placed along this continuum according to the length of the SI: <25% = 1.2, 25–49% = 1.4, 50–74% = 1.6, and 75–99% = 1.8.

**Figure 4.**
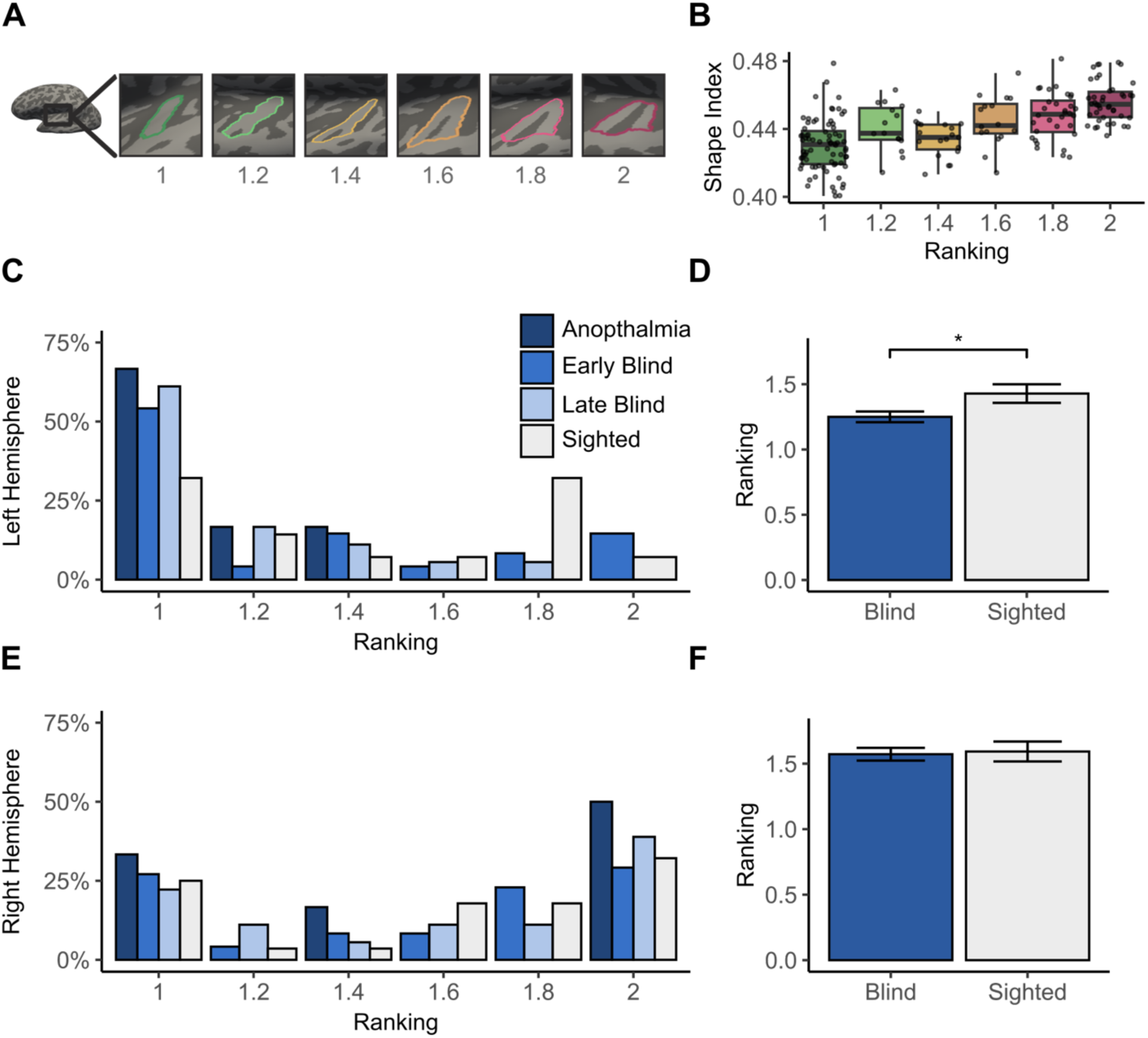
Heschl’s gyrus (HG) folding across blindness groups, categorized by the ranking method. (A) Examples of HG folding across ranks: 1 (single), 1.2 (<25% duplication), 1.4 (25–49%), 1.6 (50–74%), 1.8 (>75%), and 2 (complete duplication). (B) Relationship between HG rank (x-axis) and shape index (y-axis), with each dot representing one hemisphere (two per participant). (C, E) Distribution of ranks across groups for the left (C) and right (E) hemispheres. (D, F) Mean HG rank for blind (blue) and sighted (white) groups in the left (D) and right (F) hemispheres. Blind participants show a lower mean rank in the left hemisphere (p = .02), with no group difference in the right hemisphere. Error bars represent ± 1 standard error of the mean.

#### Bilateral categorization

Additionally, following a prior work (Benner et al., 2017), we derived a bilateral measure of HG morphology by combining HG classifications from both hemispheres. Each participant was assigned one category reflecting the folding pattern across hemispheres. Partial and complete duplications were grouped into one “duplication” category, resulting in four categories in total (left/right hemispheres): single/single, single/duplication, duplication/single, and duplication/duplication (**Figure 5A**). For this analysis, we applied the classification scheme from Da Costa et al (2011).

**Figure 5.**
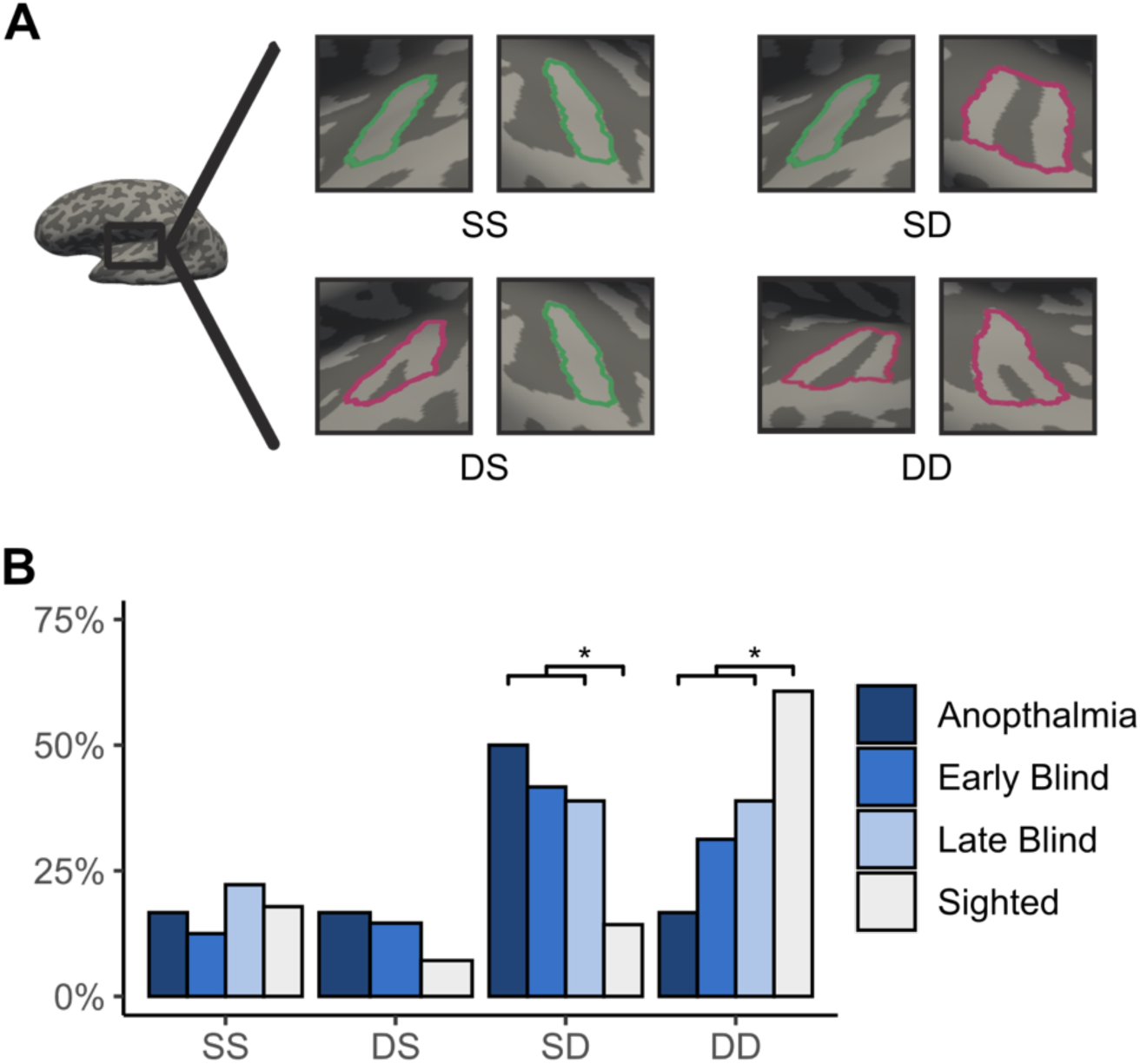
Heschl’s gyrus (HG) folding across blindness groups with joint categorization. (A) Examples of bilateral HG configurations: SS (single in both hemispheres), DS (duplication left, single right), SD (single left, duplication right), and DD (duplication in both hemispheres). (B) Percentage of each configuration across groups (anophthalmia, early blind, late blind, sighted controls). Blind individuals show a higher prevalence of SD and a lower prevalence of DD compared to sighted controls (p = .05).

### Continuous Cortical Folding and Other Structural Measurements

Alongside the visual categorization of folding, we computed a set of continuous surface-based metrics to quantify the amount of cortical folding in HG. We used FreeSurfer’s `mris_anatomical_stats` to extract the following metrics: mean cortical thickness, total surface area, mean gray matter volume, mean Curvedness Index, and mean Shape Index (Fischl, 2012; Koenderink & van Doorn, 1992a). Additionally, we calculated the local Gyrification Index (Schaer et al., 2008) by averaging vertex-wise values within the HG ROI. All metrics were calculated separately for each hemisphere and each HG ROI.

### Experimental Design and Statistical Analysis

All statistics were computed with R-4.4.1 in RStudio (Posit team, 2025; R Core Team, 2024; Wickham et al., 2019). Chi-square tests of independence were conducted to compare the distribution of folding categorizations across blindness groups. We ran Mann-Whitney U tests to compare the mean rankings across groups and Wilcoxon signed-rank tests to compare the mean rankings across hemispheres. Lastly, we performed a linear mixed-effects model for the continuous metrics. This model treated age, group, and hemisphere as fixed effects, and MRI scan location and participant (nested) as random effects:

Y ∼ age + hemisphere + group + random effects(location:participant).

For all chi-square and non-parametric tests, we combined our participant groups in two ways given the sample sizes per group. First, all blind individuals were combined into a single group regardless of blindness onset to consider the absence or presence of vision (ANO/EB/LB vs. SC). Second, participants were grouped based on auditory experience during early development (ANO/EB vs. LB/SC), allowing us to test whether early auditory experience during periods of heightened developmental sensitivity shapes HG morphology. For all linear mixed-effects models, blind groups were separated by onset: ANO, EB, and LB. Five late blind participants from Jiang et al. (2016) and nine sighted control participants from the Oxford University dataset were excluded from the linear mixed-effects models due to missing demographic information.

## Results

### Da Costa et al. method

We began by classifying the HG morphology in our participant groups for each hemisphere using the method from Da Costa et al. (2011). The method relies on surface-based morphology and categorizes an HG shape with any splits united by a common stem as a partial duplication (**Figure 2A**). The distribution of single, partial, and complete duplications in each hemisphere for each group is summarized in **Figures 2B and 2C**.

To validate our classification approach, we first compared the HG morphology distribution in our sighted group with that reported by Da Costa et al. The prevalence of each morphology type in our study was comparable to their findings (left hemisphere: 𝜒^2^(2, N = 28) = 1.29, *p* = 0.53; right hemisphere: 𝜒^2^(2, N = 28) = 0.96, *p* = 0.62), supporting the consistency of our method. Also consistent with previous findings (Abdul-Kareem & Sluming, 2008; Campain & Minckler, 1976), we observed that, for our sighted control group, complete duplications were more common in the right hemisphere (32%) than in the left hemisphere (7%), and a partial duplication (52%) was more frequently observed than a complete duplication (20%) across both hemispheres. These confirm that our classification replicates established morphological distributions in sighted individuals, providing a reliable baseline for comparisons with blind participants.

Then, to answer our main question, we examined HG morphology in blind individuals and compared it with that of our sighted controls. On average, across all the blind groups, we observed a similar pattern of HG morphology as observed in sighted individuals: complete duplications were also more common in the right hemisphere (33%) than the left hemisphere (10%), and partial duplications (37%) were more common than complete duplications (22%) across both hemispheres.

A chi-squared analysis revealed that our blind groups did not have a higher frequency of duplications than the sighted controls under the blindness grouping (ANO/EB/LB vs. SC). HG morphology types were not significantly different between the sighted and blind participants in the right hemisphere (𝜒^2^ (2, N = 100) = 0.06, *p* = 0.97). In the left hemisphere, results trended in the *opposite* direction of our predicted pattern of results (𝜒^2^ (2, N = 100) = 6.32, *p* = 0.04) where a post-hoc test revealed the blind groups tended to have a higher frequency of singles and lower frequency of partials in the left hemisphere than the sighted group, although this was not statistically significant (*p* = .08, multiple-comparison corrected).

No significant effects were observed under the developmental grouping (ANO/EB vs. LB/SC) in either hemisphere (left hemisphere: 𝜒^2^ (2, N = 100) = 5.38, *p* = 0.07; right hemisphere: 𝜒^2^ (2, N = 100) = 0.23, *p* = 0.90), indicating that HG morphology did not vary as a function of early auditory experience.

### Marie et al. method

Next, we examined the effects of blindness on the HG morphology types using the method from Marie et al.’s study—one of the studies with the largest sample of normally sighted individuals to characterize HG morphology type using MRI (N = 430, 232 right-handers). Their method primarily used a volumetric approach where a single sagittal volume was chosen to assess the presence of partial or complete duplications within HG. This study considers an HG to be a partial if there is an SI at least 1/3 the length of the HG. Given this method, any partials with a split less than 1/3 of the size of the HG would be considered a single (**Figure 3A**).

Using this approach, within our sighted controls, we again found that the prevalence of HG morphology types was statistically comparable to that reported by Marie et al. findings (left hemisphere: 𝜒^2^(2, N = 28) = 2.98, *p* =0.23; right hemisphere: 𝜒^2^(2, N = 28) = 4.14, *p* = 0.13), giving us high confidence in our manual labeling.

Consistent with the Da Costa et al. method, we found no evidence that our blind groups had a higher probability of duplications in either hemisphere than sighted controls (**Figures 3B & 3C**). Under both grouping schemes, there was no effect of group in the right hemisphere (blindness grouping: 𝜒^2^ (2, N = 100) = 0.68, *p* = 0.71; developmental grouping: 𝜒^2^ (2, N = 100) = 0.20, *p* = 0.90).

In the left hemisphere, we again observed a significant effect under the blindness grouping in the *opposite* direction of our prediction (𝜒^2^ (2, N = 100) = 10.41, *p* = 0.01). Post-hoc pairwise comparisons indicated that the combined blind group showed a higher proportion of single gyri and fewer partial duplications than the sighted controls (*p* = .01, multiple-comparison corrected). A similar overall effect was observed under the developmental grouping (𝜒^2^ (2, N = 100) = 6.14, *p* = 0.01); however, no post-hoc tests were significant when adjusting for multiple comparisons.

### Graded duplication ranking method

Finally, we developed a novel classification system that ranked HG by the degree of duplication. Visual depictions of the six ranks are in **Figure 4A**. To validate this approach, we compared our rankings with the Shape Index, a well-established quantitative characterization of local surface shape along a continuum from sulcal (concave) to gyral (convex; Koenderink & van Doorn, 1992b). We expected higher duplication ranks to correspond to greater Shape Index values because more duplications of HG should correspond with greater local concavity. Performing a Spearman correlation revealed our rankings and Shape Index are positively correlated (*r*_S_(198) = 0.60, *p* < 0.001; **Figure 4B**), supporting the validity of our method.

**Figures 4C-D** show the folding patterns observed in the blind and sighted groups using the graded ranking method. An overall hemispheric asymmetry was again observed, consistent with previous results (Campain & Minckler, 1976). For the sighted controls, a paired Wilcoxon signed-rank test revealed that the mean duplication rank in the right hemisphere (1.59) was higher than the left hemisphere (1.43; V = 21, *p* = 0.03), demonstrating an overall higher degree of duplication in the right hemisphere. Similarly, a higher duplication ranking in the right hemisphere than the left was also observed in the blind group (left = 1.25; right = 1.57; V = 249.5, *p* < 0.001).

Turning to our main question, we again found no overall effect of blindness in the right hemisphere. Under both grouping schemes, duplication rank did not differ across groups in the right hemisphere (blindness grouping: U = 995, *p* = 0.92; developmental grouping: U = 1198, *p* = 0.76).

In the left hemisphere, consistent with our observations using established categorization methods, the blind group showed a significantly lower mean rank (i.e., less duplication) than the sighted controls (U = 735, *p* = 0.02). However, this effect was not observed under the developmental grouping (U = 1138.5, *p* = 0.44).

### Bilateral categorization

Contrary to our initial hypothesis, blind individuals tend to have a higher proportion of single HGs and fewer duplications than sighted controls. Notably, this effect was observed only in the left hemisphere, while the right hemisphere showed no group difference.

Next, to examine if this effect depends on bilateral folding patterns, we assigned each participant to a category representing the combined HG morphology pattern across hemispheres **(Figure 5A)**. In the general population, the two most prevalent configurations are bilateral duplication, and single in the left hemisphere with duplication on the right (Campain & Minckler, 1976). Among our sighted participants, the most common folding pattern was the first of these two configurations: duplication in both hemispheres (62%). Blind individuals tended towards the other prevalent configuration: single on the left and duplication on the right (42%). A duplication in the left hemisphere and a single in the right hemisphere was the least common pattern for both sighted (7%) and blind (11%) individuals.

Accordingly, a chi-squared test revealed a significant difference in bilateral morphology categories between blind and sighted individuals under the blindness grouping (𝜒^2^ (3, N = 100) = 9.02, *p* = 0.03). Specifically, among participants with a duplication in the right hemisphere, blind individuals were more likely to have a single in the left hemisphere than sighted individuals (*p* = .05, multiple-comparison corrected; **Figure 5B**). A similar pattern was observed under the developmental grouping (𝜒^2^ (3, N = 100) = 9.10, *p* = 0.03). Individuals who were blind early during development (ANO and EB) were more likely to show a single HG in the left hemisphere when the right hemisphere was duplicated, compared to those with visual experience during early development (LB and SC; *p* = .03, multiple-comparison corrected).

### Continuous cortical folding measurements

In addition to the manual categorization of HG morphology, we examined two continuous measures of cortical folding: the Gyrification Index and the Curvedness Index (Koenderink & van Doorn, 1992a; Schaer et al., 2008). Whereas manual labeling captures gross folding shapes within HG, these metrics provide complementary, quantitative descriptions of geometry at a continuous scale. Specifically, the Gyrification Index quantifies the ratio of the total cortical surface area to the exposed outer surface, indexing the overall degree of cortical folding. The Curvedness Index measures the magnitude of local curvature at each surface vertex, describing how sharply the cortex bends regardless of whether it is convex or concave.

In our sighted participants, the Gyrification Index in HG showed a negative relationship with age (*r*(36) = -0.63, *p* < 0.001), indicating that overall gyrification decreased with age (**Figure 6A**). This negative relationship has been reported globally across brain regions (Hogstrom et al., 2013). Given this strong relationship with age and the hemispheric asymmetries observed in folding categories, we used a linear mixed-effects model to test for the effects of blindness while controlling for these factors. The models included group (ANO, EB, LB, and SC), hemisphere, and age as fixed effects and scan location and participant (nested) as random effects.

**Figure 6.**
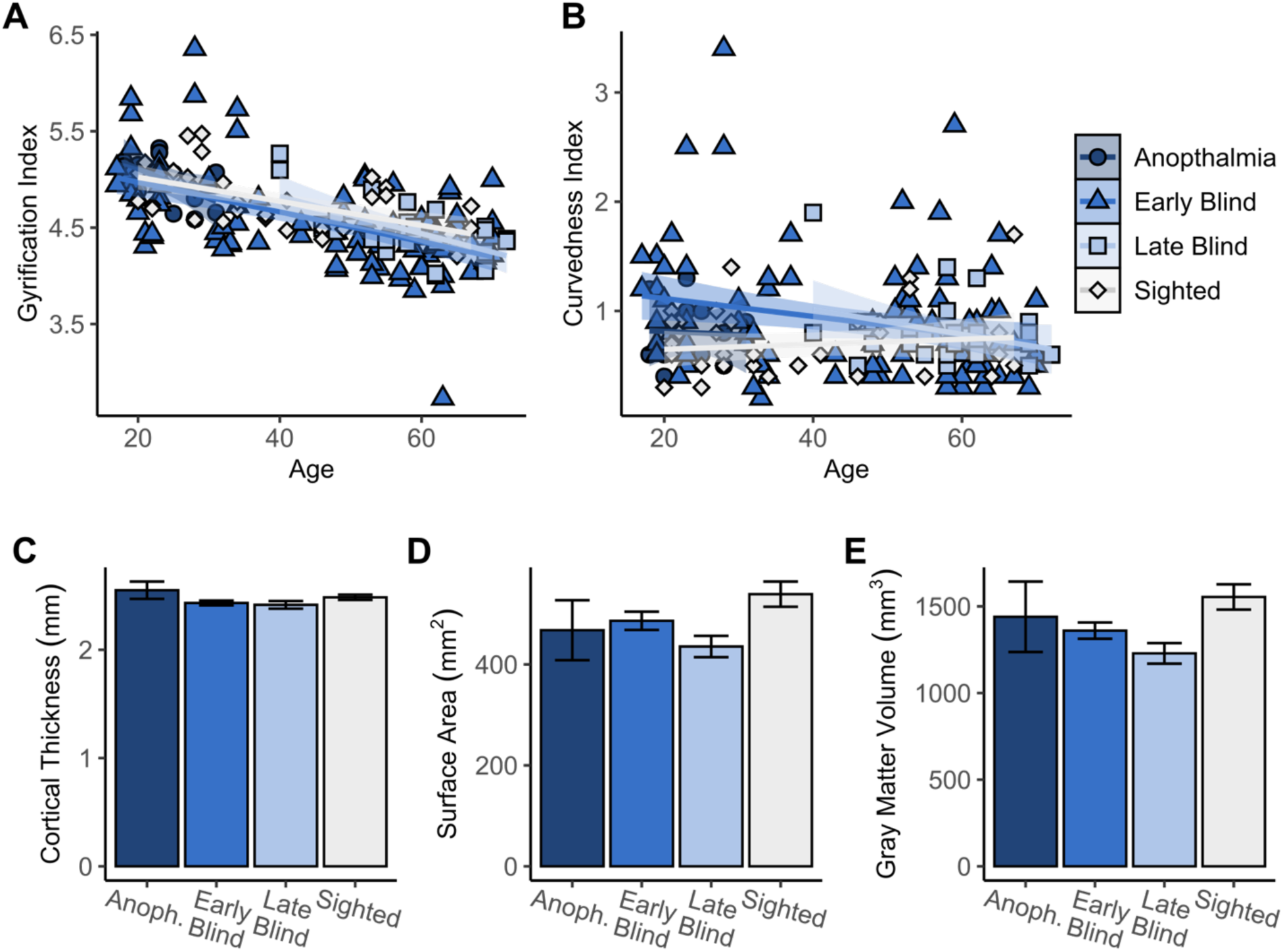
Gyrification Index, Curvedness Index, cortical thickness, surface area, and gray matter volume across blindness groups. (A–B) Mean Gyrification and Curvedness Indices of HG as a function of age, with best-fit lines for each group. Each dot represents a hemisphere (two per participant); symbols denote group (anophthalmia: circle; early blind: triangle; late blind: square; sighted: diamond). (C–E) Mean cortical thickness (mm), surface area (mm*²*), and gray matter volume (mm*³*) of HG across groups. Color shading (light to dark blue) indicates late blind, early blind, and anophthalmic participants; sighted controls are shown in white. Error bars represent ±1 standard error of the mean.

If extensive auditory experience increases HG folding, we should see higher Gyrification Index and Curvedness Index in our blind groups than in the sighted group. However, contrary to our predictions, the linear mixed-effects model revealed no significant main effect of group on Gyrification Index (F(3,81) = 0.61, *p* = .61, **Figure 6A**) nor the Curvedness Index (*F*(3, 4.48) = 0.48, *p* = .71, **Figure 6B**).

### Other structural metrics

Given prior reports of experience-related changes in other structural metrics within the left hemisphere HG (Golestani et al., 2011; Schneider et al., 2005; Wong et al., 2008), we explored additional cortical structural measures, including cortical thickness, surface area, and gray matter volume (see **Figure 6**).

Using the same linear mixed-effects models as above, we found no significant group differences in any of these metrics (**Figures 6C-E**; thickness: *F*(3, 5.88) = 0.75, *p* = 0.56; surface area: F(3, 2) = 0.81, p = 0.59; gray matter volume: F(3, 81) = 0.99, p = 0.40). There was an effect of hemisphere in the cortical thickness and surface area, but only trending in volume, where all measures were higher in the right hemisphere than the left hemisphere (thickness: F(1, 85) = 4.57, p = 0.04; surface area: F(1, 85) = 4.27, p = 0.04; gray matter volume: F(1, 85) = 3.59, p = 0.06). These results further suggest that auditory experience associated with blindness does not produce structural changes in HG. Full model results are provided in **Table 2**.

**Table 2.**
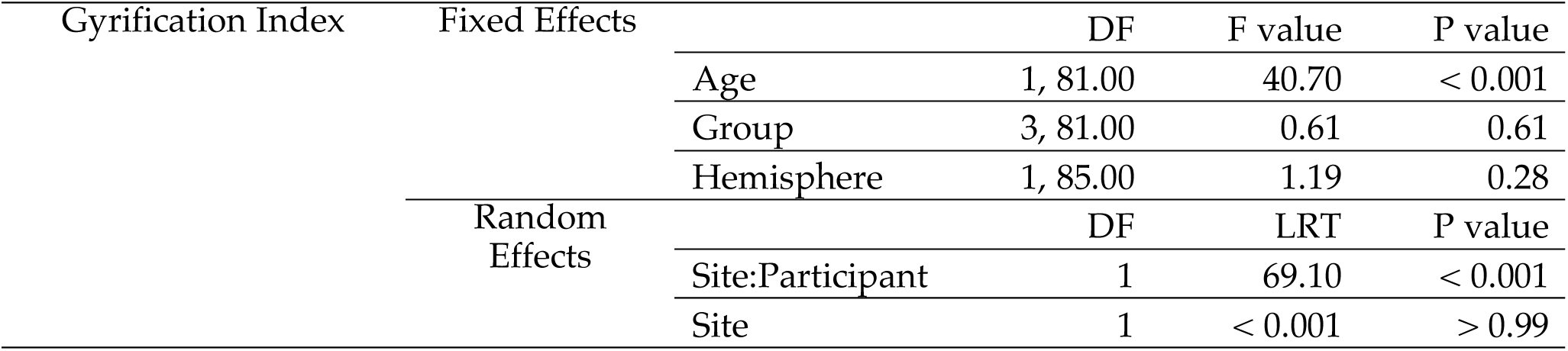
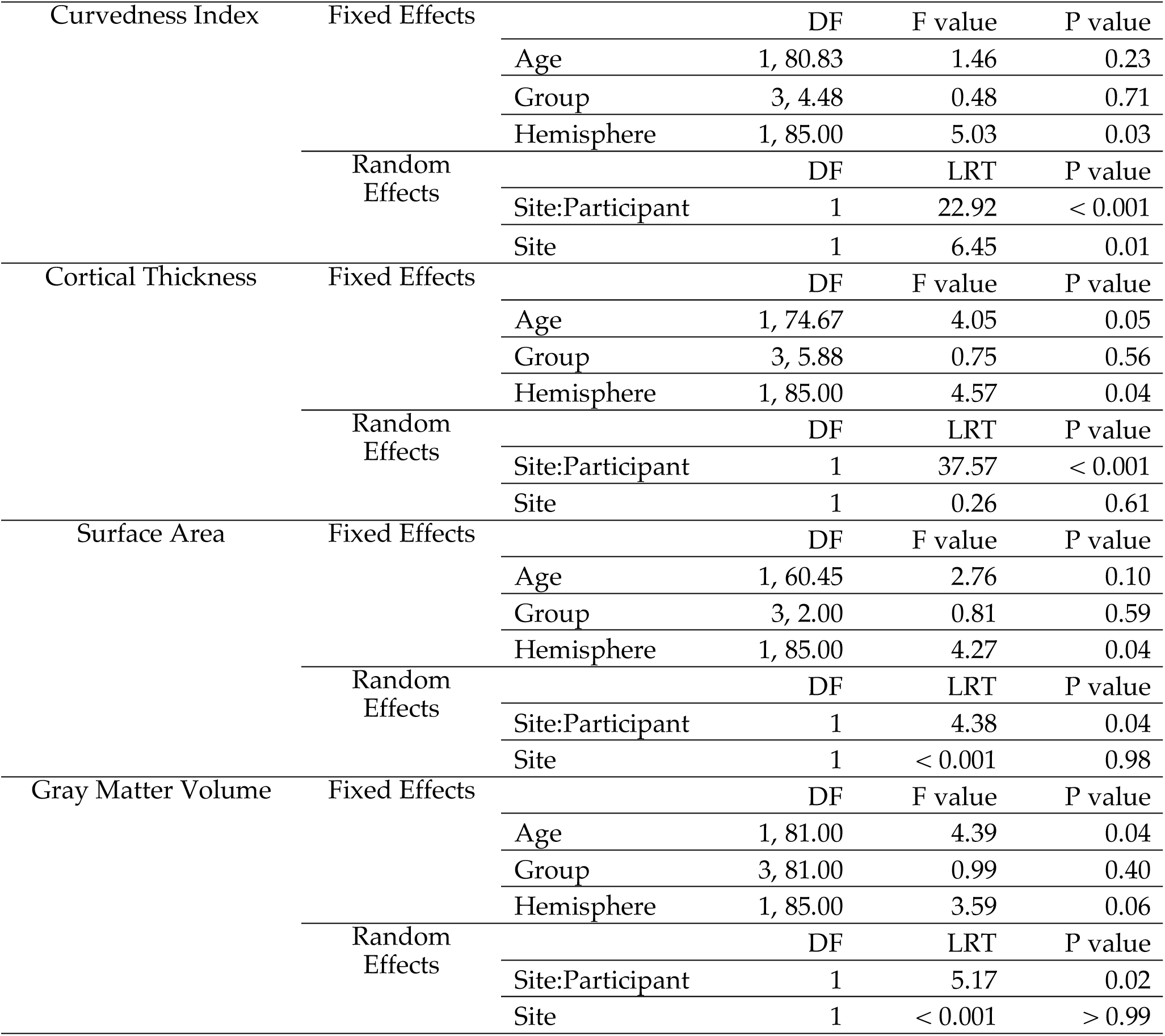
Linear mixed-effect model results for quantitative measures: Gyrification Index, Curvedness Index, cortical thickness, surface area, and gray matter volume. For all measures, group, hemisphere, and age were fixed effects. Participant and scan site were nested as random effects. This table lists the degrees of freedom (DF, if applicable; numerator, denominator), the test statistic (either the F value or the Likelihood Ratio Test (LRT)), and the P value. Values are rounded to two decimal places.

| Gyrification Index | Fixed Effects | DF | F value | P value |
| --- | --- | --- | --- | --- |
|  | Age | 1, 81.00 | 40.70 | < 0.001 |
|  | Group | 3, 81.00 | 0.61 | 0.61 |
|  | Hemisphere | 1, 85.00 | 1.19 | 0.28 |
|  | Random Effects | DF | LRT | P value |
|  | Site:Participant | 1 | 69.10 | < 0.001 |
|  | Site | 1 | < 0.001 | > 0.99 |

**Table 2. Linear mixed-effect model results for quantitative measures: Gyrification Index, Curvedness Index, cortical thickness, surface area, and gray matter volume.**
|  |  |  |  |  |  |
| --- | --- | --- | --- | --- | --- |
| Curvedness Index | Fixed Effects |  | DF | F value | P value |
|  |  | Age | 1, 80.83 | 1.46 | 0.23 |
|  |  | Group | 3, 4.48 | 0.48 | 0.71 |
|  |  | Hemisphere | 1, 85.00 | 5.03 | 0.03 |
|  | Random Effects |  | DF | LRT | P value |
|  |  | Site:Participant | 1 | 22.92 | < 0.001 |
|  |  | Site | 1 | 6.45 | 0.01 |
| Cortical Thickness | Fixed Effects |  | DF | F value | P value |
|  |  | Age | 1, 74.67 | 4.05 | 0.05 |
|  |  | Group | 3, 5.88 | 0.75 | 0.56 |
|  |  | Hemisphere | 1, 85.00 | 4.57 | 0.04 |
|  | Random Effects |  | DF | LRT | P value |
|  |  | Site:Participant | 1 | 37.57 | < 0.001 |
|  |  | Site | 1 | 0.26 | 0.61 |
| Surface Area | Fixed Effects |  | DF | F value | P value |
|  |  | Age | 1, 60.45 | 2.76 | 0.10 |
|  |  | Group | 3, 2.00 | 0.81 | 0.59 |
|  |  | Hemisphere | 1, 85.00 | 4.27 | 0.04 |
|  | Random Effects |  | DF | LRT | P value |
|  |  | Site:Participant | 1 | 4.38 | 0.04 |
|  |  | Site | 1 | < 0.001 | 0.98 |
| Gray Matter Volume | Fixed Effects |  | DF | F value | P value |
|  |  | Age | 1, 81.00 | 4.39 | 0.04 |
|  |  | Group | 3, 81.00 | 0.99 | 0.40 |
|  |  | Hemisphere | 1, 85.00 | 3.59 | 0.06 |
|  | Random Effects |  | DF | LRT | P value |
|  |  | Site:Participant | 1 | 5.17 | 0.02 |
|  |  | Site | 1 | < 0.001 | > 0.99 |

## Discussion

Here, we examined whether blindness alters cortical folding in HG, motivated by the idea that extensive auditory experience might lead to increased folding within the auditory cortex. Contrary to this prediction, we found no evidence that blindness increases HG folding. Across multiple approaches, including morphological classifications and continuous surface-based measures, HG folding was not increased in blind individuals compared to sighted controls. Importantly, these analyses were conducted in the largest dataset of blind participants examined to date (N = 72 across groups).

### Blindness does not increase cortical folding in HG

We addressed variability in how HG folding has been labeled across prior studies by applying several established classification schemes and by developing a new ranking system that orders HGs by degree of duplication. In addition to these manual gold-standard approaches, we also examined continuous quantitative metrics. Across all approaches, we consistently found no evidence that blindness is associated with increased HG folding, indicating that extensive auditory experience does not measurably alter HG gyrification. Our findings, therefore, are more consistent with the view that the morphology of HG is biologically predetermined and largely stable once established.

In manual labeling analyses, blind participants showed a consistent pattern characterized by a higher prevalence of single HGs in the left hemisphere. This pattern likely reflects normative variability in HG morphology rather than a blindness-specific effect on cortical folding. A single HG in the left hemisphere is among the most common folding configurations in the general population (Abdul-Kareem & Sluming, 2008), and the bilateral combination of a single left and duplicated right HG has been reported as a frequent pattern of variation as well (Campain & Minckler, 1976).

More generally, HG morphological labels may not reflect the overall degree of cortical folding. Morphological labels such as “single” or “duplication” describe how sulcal boundaries partition the gyrus but do not necessarily correspond to differences in the amount of cortical surface folded within HG. Indeed, the absence of group differences in metrics such as the Gyrification and Curvedness Indices supports this view, suggesting that while the morphological organization of HG may vary, the total cortical territory devoted to this region remains comparable between blind and sighted participants.

### Blindness does not alter cortical thickness, surface area, and gray matter volume of HG

An exploration of additional structural markers revealed no effect of blindness on cortical thickness, surface area, or gray matter volume within HG. This null result diverges from findings in the visual cortex of blind individuals, where structural alterations are observed. Specifically, studies have found increased cortical thickness and reduced surface area in the deprived occipital cortex of blind animals and humans as compared to sighted controls (Bridge et al., 2009; Dehay et al., 1991; J. Jiang et al., 2009; H.-J. Park et al., 2009). Additionally, visual deprivation has been reported to reduce cortical folding in the primary visual cortex of ferrets (Andelin et al., 2019). These findings have been interpreted as consequences of early visual deprivation disrupting the typical maturation of cortical expansion and synaptic pruning during early development, resulting in a thicker but less expanded cortex. In contrast, the auditory cortex in blindness is not deprived but remains highly active and functionally engaged. The absence of structural alterations in HG as a result of blindness suggests that deprivation and enhanced experience have differential effects on cortical development.

### Nature vs. nurture in the auditory cortex

Previous studies of auditory expertise in phoneticians and musicians have reported increased cortical thickness and gray matter volume in auditory regions, together with a greater prevalence of HG duplications (Benner et al., 2017; Golestani et al., 2011). Why is it that blindness, which is associated with lifelong extensive auditory expertise, does not produce comparable structural effects?

Numerous behavioral studies have established that blind individuals, especially those who became blind early in life, are auditory experts across many perceptual domains that overlap with those trained as musicians and phoneticians, including pitch discrimination (Gougoux et al., 2004), sound localization (F. Jiang et al., 2014; W. J. Park & Fine, 2023), and voice or speech processing (Röder et al., 2001; Stevens & Weaver, 2005). Thus, differences in the quality or degree of auditory experience per se are less likely to explain the absence of structural differences in HG.

The most plausible explanation is that previously reported structural effects in auditory expertise studies may reflect biases in career choice and career success: individuals with larger auditory cortical capacity are predisposed to pursue and excel in auditory domains. For example, in musicians, gray matter volume has been shown to vary as a function of years of musical performance in regions such as the pars opercularis (POP), which are thought to support functions related to both language and music (Abdul-Kareem et al., 2011). While this finding that career length is correlated with gray matter volume is often interpreted as evidence of experience-dependent structural plasticity, orchestral music is a highly competitive career, so it is difficult to rule out the possibility that individuals with advantageous pre-existing structural differences are more likely to have long term career success.

It is also possible that commonly used anatomical metrics, such as gray matter volume, may reflect changes in other tissue properties that are more sensitive to experience. Although one longitudinal study reported anatomical expansion within the auditory cortex following 15 months of musical training in children (Hyde et al., 2009), this work relied on voxel-based Jacobian deformation analyses, which are sensitive to shifts in tissue boundaries. As a result, the observed expansion within these trained children may instead reflect changes in tissue contrast driven by increased white matter myelination—a process that continues well into adulthood (Bethlehem et al., 2022; Paus et al., 2001). Such myelination can shift the gray–white matter boundary and alter local deformation estimates, potentially producing apparent expansion. Alternatively, early training may increase the rate of expansion of auditory cortex, without changing the final endpoint morphology.

In this context, the absence of anatomical differences in HG associated with blindness suggests that extensive auditory experience alone may not be sufficient to drive measurable anatomical changes in auditory cortex in adulthood.

This interpretation is further supported by theories of cortical development that emphasize early constraints on large-scale anatomy: folding patterns are thought to arise from mechanical and physical forces operating during early brain development, such as differential expansion between cortical and subcortical layers (Garcia et al., 2018; Tallinen et al., 2016). At the same time, while the structural geometry is likely fixed early, finer-grained functional refinement can still occur through experience (W. J. Park & Fine, 2020). Consistent with this dissociation between structure and function, recent population receptive field analyses show enhanced functional selectivity in primary and secondary auditory cortices of early blind individuals (Huber et al., 2019). Thus, it is likely that experience-dependent changes in the auditory cortex in blind individuals are primarily expressed at the level of neural tuning rather than in anatomical structure.

In conclusion, across multiple complementary analyses, we found no evidence that blindness increases HG folding. The findings suggest that the gross morphology of the HG is likely constrained by biological predispositions established early in development. Future studies that combine longitudinal imaging with microstructural and functional measures will be essential to delineate how multiple levels of plasticity interact to support auditory specialization within HG.

## Acknowledgments

NIH R01 EY014645 to IF; NIH R00 EY034546 and Georgia Institute of Technology Smithgall-Watts Early Career Award to WP.

